# Mouse higher visual areas provide both distributed and discrete contributions to visually guided behaviors

**DOI:** 10.1101/2020.03.21.001446

**Authors:** Miaomiao Jin, Lindsey L. Glickfeld

## Abstract

Cortical parallel processing streams segregate many diverse features of a sensory scene. However, some features are distributed across streams, begging the question of whether and how such distributed representations contribute to perception. We determined the necessity of primary visual cortex (V1) and three key higher visual areas (LM, AL and PM) for perception of orientation and contrast, two features that are robustly encoded across all four areas. Suppressing V1, LM or AL decreased sensitivity for both orientation discrimination and contrast detection, consistent with a role for these areas in sensory perception. In comparison, suppressing PM selectively increased false alarm rates during contrast detection, without any effect on orientation discrimination. This effect was not retinotopically-specific, suggesting a distinct role for PM in the regulation of noise during decision-making. Thus, we find that distributed representations in the visual system can nonetheless support specialized perceptual roles for higher visual cortical areas.

## Introduction

Parallel processing is a fundamental organizing principle in sensory circuits [1–4]. The visual system is particularly parallelized with a large number of higher order cortical areas that each encode overlapping portions of the visual field [5–7]. This architecture is thought to enable functional specialization and read-out of receptive fields for behavior [8,9]. However, while there are some sensory stimulus properties that seem to be both functionally and perceptually localized to specific areas [10–12], there are also a number of examples of stimulus properties that have distributed representation across the visual system [13–15]. For instance, in the mouse visual system, the sensitivity of neurons to orientation and contrast is remarkably similar across higher order cortical areas [16–19]. These distributed representations beg the question of which areas are actually responsible for perception of these features.

One possibility is that the redundancy makes it such that no single higher area is necessary for the perception of distributed stimuli. This is particularly plausible in the mouse visual system where the primary visual cortex (V1) innervates at least nine higher visual areas (HVAs; [20]). Thus, this highly parallel organization could support a robust system in which the removal of any one area has little effect on the percept. In addition, V1 also encodes simple representations such as orientation and contrast and projects to a variety of subcortical areas such as the superior colliculus and pontine nuclei [21–23]. Thus, there is a direct route from sensory stimulus to action that could bypass higher cortical areas in the execution of simple behaviors.

Another possibility is that the entire distributed network is required to enable maximal sensitivity in perceptual performance. This might happen if the readout is monitoring the output of all areas, or if there is a hierarchical structure such that each area could act as a bottleneck [24,25].

There may also be intermediate conditions, potentially reflecting specialized anatomy or readout, in which only a subset of the distributed network contributes to perceptual sensitivity. Further, different areas may make distinct contributions to task performance. Indeed, HVAs that belong to the posterior parietal cortex seem to be more important for decision-making than sensory integration, despite receiving direct input from V1 [26–28]. Thus, careful dissection of multiple features of task performance, including hit, false alarm and lapse rates, is necessary to understand the function of each area in a distributed network.

We found evidence for both distributed and discrete contributions of three HVAs (LM: lateromedial; AL: anterolateral; PM: posteromedial) to perception of broadly represented features: orientation and contrast. Like V1, AL and LM were also required for perception of both orientation and contrast. However, PM was not required for the orientation discrimination task, despite task stimulus information being robustly encoded in all areas. The most reliable effect of suppressing PM was an increase in false alarm rate during the contrast detection task. This effect was independent of the alignment of the visual stimulus with the affected cortical representation, suggesting a more global role for PM in tasks involving the detection of weak signals on a noisy background. Consistent with this hypothesis, suppression of PM increased false alarm rates during a speed increment detection task that also includes the need to overcome noise in the stimulus. Thus, our results provide evidence that distributed representations are neither completely robust to perturbation of single areas nor homogenously supported by all areas, thereby revealing clear functional specialization in the visual system.

## Results

### Rapid and reversible silencing of mouse visual cortical areas

To determine the involvement of visual cortical areas in perceptual decision-making, we suppressed neuronal activity within specific HVAs by expressing excitatory opsins (Channelrhodpsin-2 or Chronos (together abbreviated as ChR)) in GABAergic interneurons [29–31]. We targeted conditional viral expression of ChR to either V1 or an HVA (LM, AL or PM; **Figures 1A** and **S1**) in lines which express Cre recombinase in inhibitory interneurons (PV∷Cre, n=10 mice; GAD∷Cre, n=2 mice). This approach enables high temporal resolution to interleave “LED” and control trials during behavioral tasks, and reasonable spatial resolution to restrict most of the suppression to the targeted HVA.

**Figure 1.**
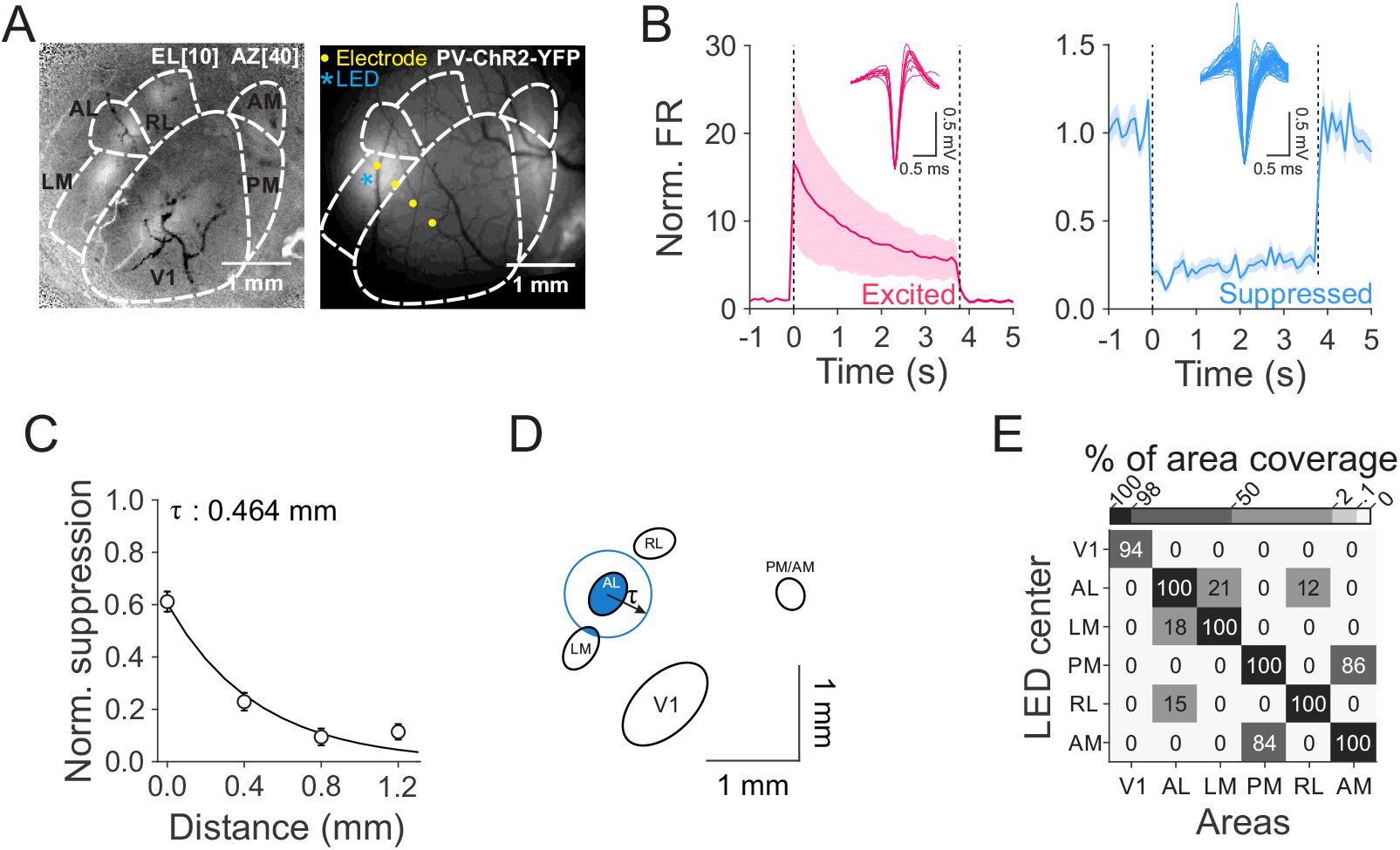
Efficacy and spatial resolution of optogenetic inhibition of mouse visual areas. (**A**) Left: change in intrinsic autofluorescence in response to a visual stimulus at elevation 10°, azimuth 40°, size 40°. Increases in fluorescence reflect activation. Right: viral expression of ChR2 in parvalbumin positive (PV+) interneurons in LM and PM. (**B**) Normalized spontaneous firing rates (FR) of cells that are excited (left, red, n=9 cells) and suppressed (right, blue, n=58 cells) when stimulated with 473 nm LED light. Dashed lines indicate light onset and offset. Insets are the waveforms of single units. ChR2 is expressed in either PV+ or GAD+ interneurons. Light power: 0.1-.5 mW across 16 experiments (n=12 mice). (**C**) Normalized suppression as a function of distance from the LED center (0 mm). Error bars are SEM across cells (n=73, 97, 81, 50 cells from 0 to 1.2 mm). The decay constant (τ) was calculated via a single exponential fit. (**D**) Oval fits for visual areas for the same example mouse in **A** with a demonstration of calculating percentage of area coverage when light is centered on AL. Blue circle was drawn based on the radius of τ from the fit in **C**. (**E**) Mean percentage of area coverage of inhibition across mice (n=15 mice).

To validate the temporal and spatial resolution, we made extracellular single unit recordings with multi-pronged electrodes that spanned up to 1.2 mm from the center of the injection site. The majority of cells near the injection site were significantly suppressed (58/82 cells; **Figure 1B, right**). Suppression was stable during the full four second duration of light stimulation (p=0.81, Kruskal-Wallis test across time bins (bin size: 100 ms), DF=36, n=58 cells, 16 experiments, 12 mice), and did not drive significant rebound excitation when the light was extinguished (first 200 ms after LED offset vs. baseline activity: p=0.40, Wilcoxon signed rank test). Thus, there is stable suppression of neuronal activity for the full duration of each behavioral trial, and independent control across trials.

We also found that spatial spread of suppression enables reasonably independent control of V1 and the HVAs. The degree of suppression rapidly decayed with distance from the injection site with a space constant of 464 μm (95% CI: [354 594 μm]; p<10^−17^, Kruskal-Wallis test, n=301 cells, **Figure 1C**). We used this space constant to estimate the degree to which suppression within any one visual area might spread to other areas. First, we measured the relative position and size of the region within V1 and each HVA that would be activated by the visual stimulus used in the behavioral tasks with intrinsic autofluorescence imaging (n=15 mice). Then, we determined the percentage of suppression of each area given its distance from the targeted area and the space constant (**Figure 1D**). This revealed that suppression targeted to V1 was well-restricted within V1 (area coverage: V1:94±2%; all other areas: 0%; **Figure 1E**), and suppression targeted to any HVA did not spread to V1. However, suppression targeted to the HVAs was not completely restricted to that HVA. For instance, while AL and LM were largely independent, there was ~20% overlap in each direction (target AL, 21±5% LM affected; target LM, 18±5% AL affected), and AM and PM share ~85% overlap (target: PM, 86±6% AM affected; target AM, 84±5% PM affected). Thus, this viral approach enables reasonable independent control of V1, LM and AL. To further improve the independent suppression of PM, we targeted our viral injection a few hundred microns posterior from the retinotopic center of PM, to avoid strong suppression of AM, but still suppress PM.

In some experiments, we also used a transgenic line to express ChR in GABAergic interneurons (VGAT-ChR2; n=2 mice; [31]). Neurons recorded in these mice exhibited clear rebound excitation after light offset (first 200 ms after LED offset vs. baseline activity: p<10^−5^, Wilcoxon signed rank test; **Figure S2A-B**). The degree of suppression was weaker (p=0.001, Wilcoxon rank sum test) and spread further away from the light center as compared to the viral approach (interaction between method and cortical distance: p<10^−6^, two-way ANOVA, **Figure S2C-E**). Therefore, we used the viral approach for the majority of experiments testing the perceptual effects of area suppression, (PV∷Cre-n=25 mice; GAD∷Cre-n=1 mouse; VGAT-ChR2-n=5 mice). Nonetheless, the behavioral results were comparable across approaches (**Figure S3**).

### Role of mouse visual cortical areas in a go/no-go orientation discrimination task

Stimulus orientation is a fundamental feature that is robustly encoded in the visual system of most species, including mouse V1 and the HVAs [18,19]. While V1 is known to be required for tasks involving discrimination of stimulus orientation [30,32–34], it is not known which, if any, higher visual areas are necessary for the perception of this distributed representation.

Therefore, we trained mice to perform a go/no-go orientation discrimination task (**Figure 2A**, [33,34]). In this task, the mice press a lever to initiate the trial and trigger the repeated presentation of a vertically oriented distractor (0°, 2-10 distractors, 100 ms duration, 250-750 ms inter-stimulus interval (ISI)) followed by the presentation of a target orientation (8-90° counter-clockwise difference from the distractor). The mice must release the lever within a brief reaction window after target onset to indicate discrimination of the target and receive water reward. Release of the lever within 200-550 ms after target onset was counted as a “hit”, while failure to release during this window was counted as a “miss”. We thus quantified hit rate as a function of target orientation to construct a psychometric function to measure discrimination threshold (**Figure 2B-E,** left). Similarly, if animals released the lever within the same window following a distractor presentation, it was categorized as a false alarm (FA), and otherwise a correct reject (CR), giving us a measure of FA rate (**Figure 2B-E,** right).

**Figure 2.**
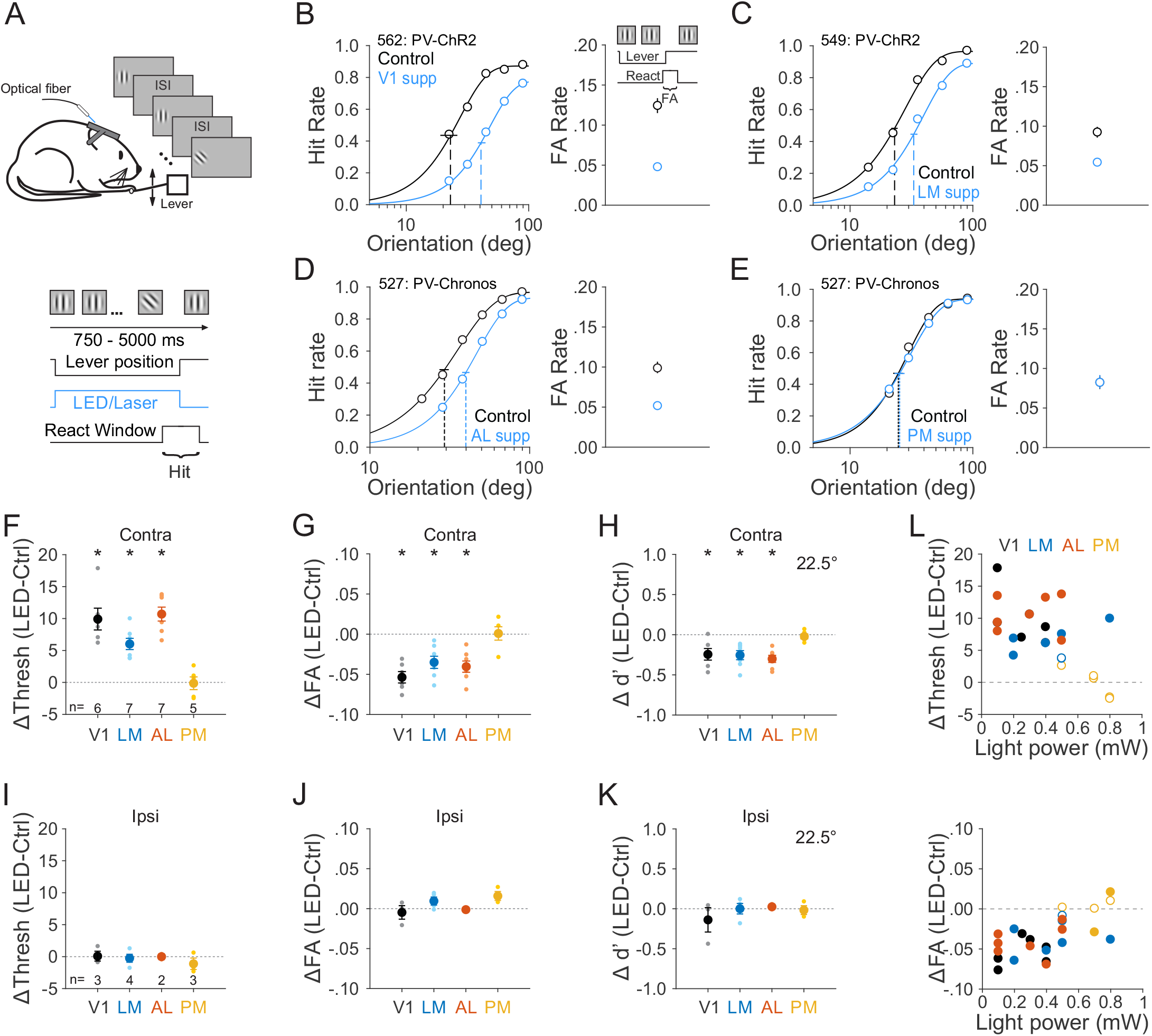
Role of mouse visual areas in discriminating orientations. (**A**) Schematic of orientation discrimination task. (**B-E**) Effect of suppression in area V1 (**B**), LM (**C**), AL (**D**), and PM (**E**) on the task performance for each representative mouse. Note that data from AL and PM come from the same mouse. Left-hit rate (Hit/(Hit + Miss)) as a function of target orientations. Data were fitted by Weibull functions to determine the thresholds (dotted vertical lines) and 95% confidence intervals (solid horizontal lines). Right-false alarm (FA) rate calculated as FA/(FA + CR). CR stands for correct rejection. Error bars are 95% confidence intervals. (**F-I**) Summary of effect of suppression in visual areas in terms of the change in the threshold (**F**), FA rate (**G**) and sensitivity (d’ for 22.5° target, **H**). Big circles are the population mean and small circles are data for each mouse. The visual stimulus is presented in the contralateral field of view (relative to the ChR2 injection site). Error bars indicate ± SEM across mice (n=6, 7, 7, 5 for V1, LM, AL, and PM respectively). * p<0.05. (**I-K**) Same as **F-H**, for visual stimulus presentation in the ipsilateral field of view. n=3, 4, 2, 3 for V1, LM, AL and PM respectively. (L) Effects of area suppression on orientation discrimination threshold (top) and FA rate (bottom) as a function of light power. Different colors denote different visual areas. Filled and open circles denote significant and non-significant difference between control and LED trials, respectively.

To determine the role of visual cortical areas on this task, we suppressed activity in the cortical hemisphere contralateral to the visual stimulus on interleaved “LED” trials, to specifically suppress visually-evoked activity. Suppression was initiated at the time of trial onset and terminated after the mouse released the lever or the reaction time expired. Therefore, we could compare task performance between control and LED trials for different visual areas (**Figures 2B-E and Supplementary item 1**). Notably, we used relatively low light powers to avoid heating the brain or impairing the animals on the easiest trials (i.e. 90° targets), so that we could avoid confounds associated with changes in task engagement.

Suppressing V1, LM and AL significantly increased the animals’ orientation discrimination threshold (V1: p=0.002, n=6 mice; LM: p<10^−3^, n=7; AL: p<10^−3^, n=7; paired t-test; **Figure 2F and 2L top**) and decreased the FA rate (V1: p<10^−3^; LM: p=0.004; AL: p=0.001; paired t-test; **Figure 2G and 2L bottom**). The magnitude of these effects was equivalent across all three areas (threshold: p=0.65; FA: p=0.22; one-way ANOVA across areas), suggesting that V1, LM and AL are similarly involved in orientation discrimination. However, when we suppressed area PM, we did not observe any significant difference in discrimination threshold (p=0.90, n=5 mice) or FA rate (p=0.92), suggesting a lack of role for PM.

This lack of effect with PM suppression cannot be explained by the efficacy of suppression since we observed similar degree of effect of ChR activation in PM as in V1 and AL, and only a slightly stronger effect in LM (Kruskal-Wallis test (p=0.007) with post-hoc Dunn's test comparing to PM (n=20 cells): V1: p=0.92, n=54; AL: p=0.09, n=28; LM: p=0.03, n=6; **Figure 3A**). The lack of effect of suppression of PM on this task was also not clearly explained by differences in visual response properties, since neurons in all areas were similarly orientation tuned (circular variance: p=0.18; V1: n=272 cells; LM: n=111 cells; AL: n=136 cells; PM: n=107 cells; One-way ANOVA; **Figure 3B**) and driven equally well by the task stimuli (comparison with PM (n=71 cells): response to 1^st^ distractor stimulus-V1: p=0.1, n=201; LM: p=0.43, n=77; AL: p=0.11, n=100; response to 4-5^th^ distractor stimuli-V1: p=0.003; LM: p=0.42; AL: p=0.12; Kruskal-Wallis test with post-hoc Dunn's test; **Figure 3C**). Thus, although task-relevant information is encoded in PM, the area is not used for discriminating stimulus orientation.

**Figure 3.**
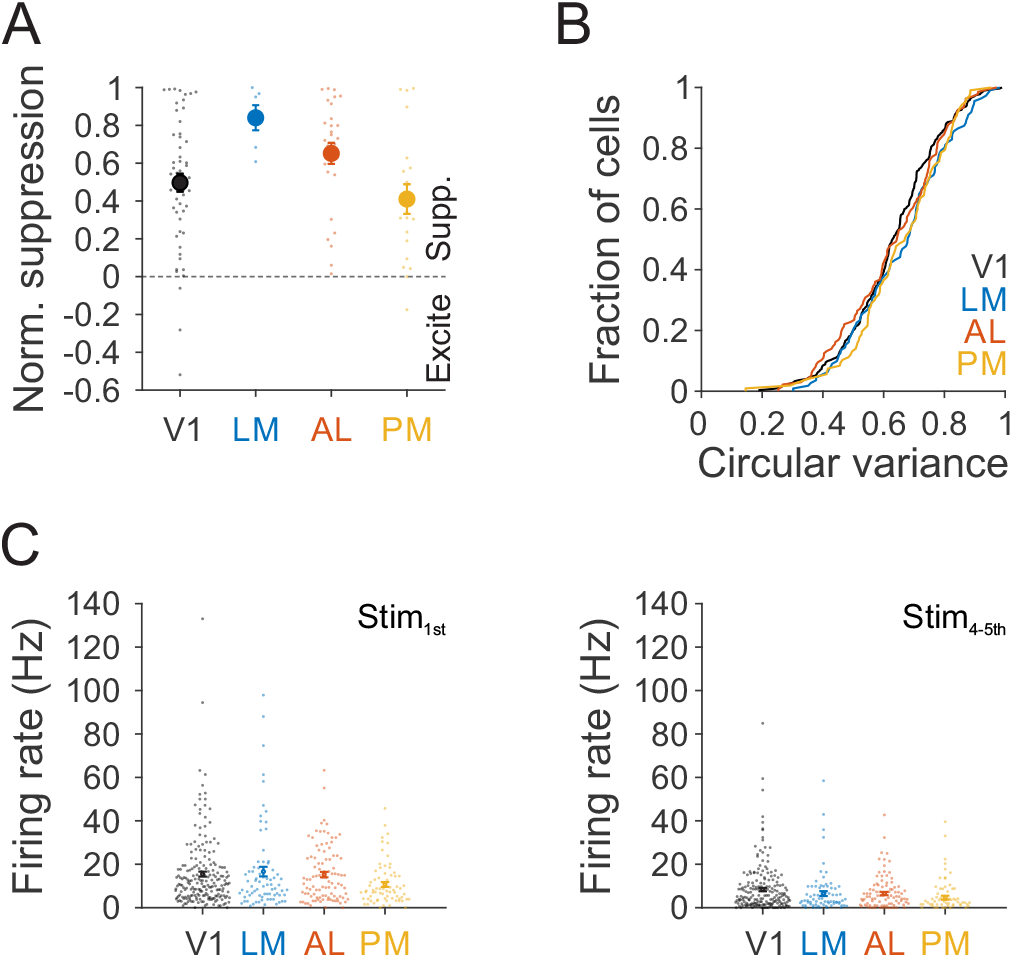
Neurons in PM have similar suppression efficacy, orientation tuning and task driven responses as V1, LM and AL. **(A)** Normalized suppression in the LED center for each area. Data from viral and transgenic approaches were combined (**Figure 1**). Small dots are individual cells, filled circles are means across population and error bars are SEM across cells (viral/transgenic: V1: n=29/25; LM: n=6/0; AL: n=17/11; n=6/14). (**B**) Cumulative distribution of circular variance of neurons' responses to different orientations across four visual areas (V1: n=272; LM: n=111; AL: n=136; PM: n=107 cells). (**C**) Neuronal firing rate in response to the first (left) and plateaued fourth and fifth distractors (right) for neurons that were significantly driven by the distractor stimulus (V1: n=201; LM: n=77; AL: n=100; PM: n=71 cells).

Despite the coincident decrease in both hit and FA rate, suppressing V1, LM and AL significantly decreased sensitivity (d’ for 22.5° target: V1: p=0.02; LM: p=0.004; AL: p<10^−3^; **Figure 2H**; [35]), consistent with a sensory role for these areas in orientation discrimination. Indeed, when we moved the visual stimulus presentation to the ipsilateral field of view (relative to the suppression site; **Supplementary item 2**), all effects of cortical suppression were abolished (threshold-V1: p=0.93, n=3 mice; LM: p=0.71, n=4; AL: p=0.90, n=2; PM: p=0.32, n=3; FA rate-V1: p=0.63; LM: p=0.16; AL: p=0.66; PM: p=0.10; sensitivity-V1: p=0.46; LM: p=0.99; AL: p=0.65; PM: p=0.79; **Figure 2I-K**). This suggests that the effects of suppression of V1, LM and AL on performance of orientation discrimination are specific and restricted to the silenced retinotopic positions.

Non-visual aspects of task performance, such as task engagement and target expectation, were not affected by suppression of the visual cortical areas (**Figure S4**). Firstly, we found no change in the lapse rate (1-hit rate at the easiest target) when suppressing V1 (p=0.33) or PM (p=0.40). We did observe a small but significant increase in lapse rate with suppression of LM (p=0.04) and AL (p=0.005, paired t-test; **Figure S4A**), however there was a strong correlation between the change in lapse rate and the change in threshold and FA rate, suggesting that this is due to a visual impairment rather than task engagement (threshold: r=0.54, p=0.006; FA rate: r=-0.36, p=0.08). Secondly, target expectation also changed over the course of a trial, such that animals reliably had lower thresholds (p<10^−5^, n=15 mice, one-way ANOVA) and higher FA rates (p<10^−6^) late in the trial, due to an increase in the probability of target occurrence with the hold time. Silencing visual areas did not affect the dependence of either threshold or FA rate on trial length (interaction between area suppression and trial length: threshold- V1: p=0.74; LM: p=0.58; AL: p=0.58; PM: p=0.58; FA rate- V1: p=0.76; LM: p=0.99; AL: p=0.84; PM: p=0.67; Two-way ANOVA; **Figure S4B-C**), suggesting a lack of a role for these areas in expectation.

In summary, we found that V1 and higher visual areas LM and AL, but not PM were required for this orientation discrimination task. The fact that the effects of V1, LM and AL suppression had no effect on non-visual aspects of the task and were specific to the affected visual field supports a mainly perceptual role of these areas.

### Role of mouse visual cortical areas in a go/no-go contrast detection task

Stimulus contrast, like orientation, robustly modulates neuronal activity across all visual areas [16,17]. To determine which, if any of the three HVAs is required for perception of this broadly represented stimulus feature, we next trained mice on a go/no-go contrast detection task ([30,36]; **Figure 4A**).

**Figure 4.**
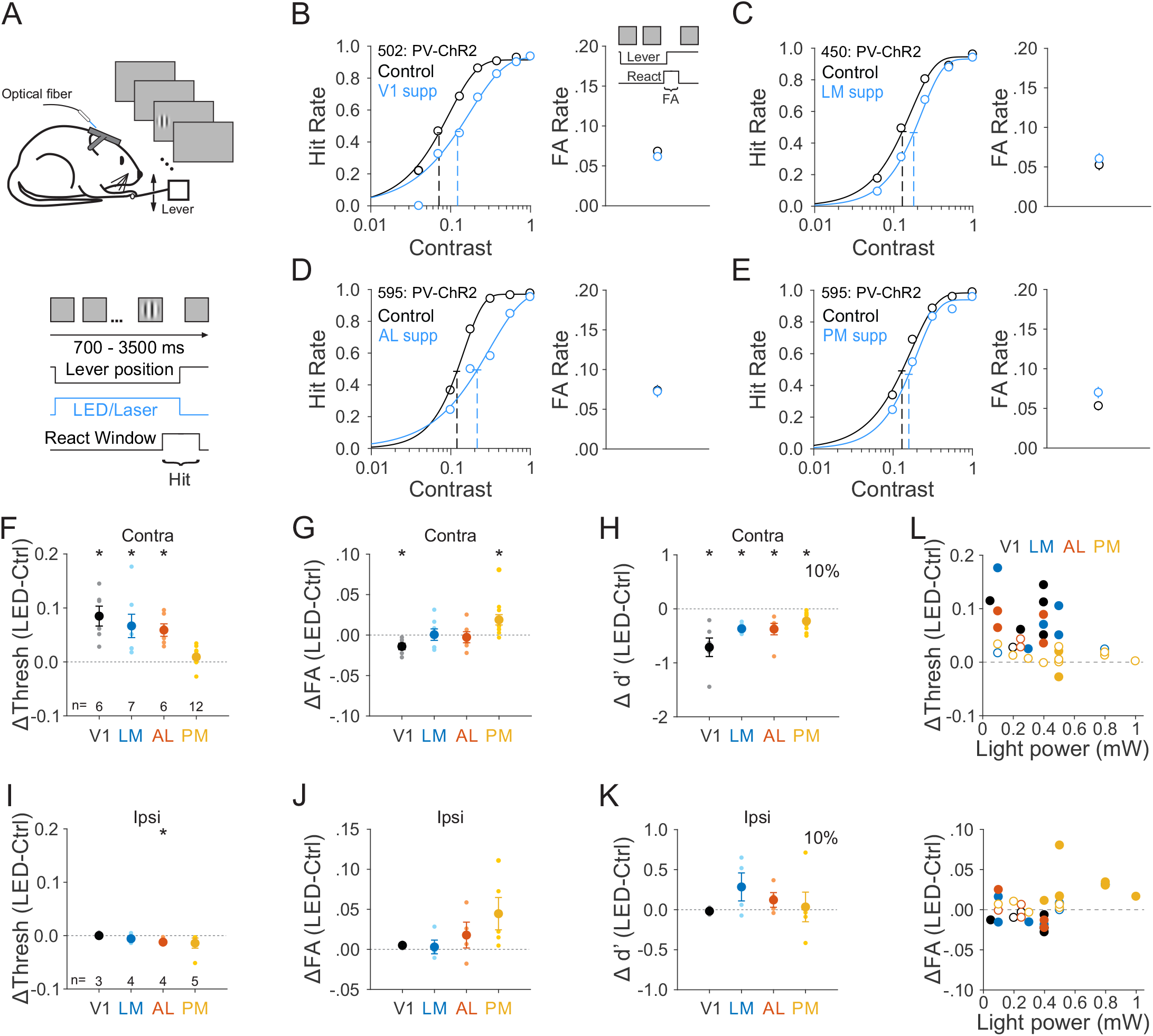
Role of mouse visual areas in detecting stimulus contrast. (**A**) Schematic of contrast detection task. (**B-E**) Effect of suppression in area V1 (**B**), LM (**C**), AL (**D**) and PM (**E**) on performance for each representative mouse. Note that data from AL and PM come from the same mouse. Left-hit rate as a function of stimulus contrast. Data were fitted by Weibull functions to determine the thresholds (dotted vertical lines) and 95% confidence intervals (solid horizontal lines). Right-FA rate. Error bars are 95% confidence intervals. (**F-H**) Summary of effect of suppression in visual areas in terms of the change in the threshold (**F**), FA rate (**G**) and sensitivity (d’ for 10% contrast, **H**). Big circles are the population mean and small circles are data for each mouse. The visual stimulus is presented in the contralateral field of view (relative to the ChR2 injection site). Error bars indicate ± SEM across mice (n=6, 7, 6, 12 for V1, LM, AL, and PM respectively). * p<0.05. (**I-K**) Same as **F-H**, for visual stimulus presentation in the ipsilateral field of view. n=3, 4, 4, 5 for V1, LM, AL and PM respectively. (**L**) Effects of area suppression on contrast detection threshold (top) and FA rate (bottom) as a function of light power. Different colors denote different visual areas. Filled and open circles denote significant and non-significant difference between control and LED trials, respectively.

As in the orientation discrimination task, in the contrast detection task the mice must press the lever to initiate each trial and release the lever to report detection of a contrasted target grating (0°, 100 ms duration, contrast range: 4%-100%; **Figure 4A**). We quantified hit rate by calculating percentage of target presentations in which the mouse released within the reaction time window (200-550 ms) after target onset (**Figure 4B, left**). We also calculated FA rate by assessing the percentage of releases within a similar window before target appearance (**Figure 4B, right**).

We found that suppressing V1, LM and AL significantly increased the contrast detection threshold (V1: p=0.006, n=6 mice; LM: p=0.02, n=7; AL: p=0.004, n=6; paired t-test; **Figure 4F, 4L top and Supplementary item 3**), while suppression of PM slightly, but not significantly, increased the threshold (p=0.08, n=12), resulting in a significantly weaker effect of suppression of PM than other areas (PM vs. V1: p<10^−4^; PM vs. LM/AL: p=0.02; one-way ANOVA (p<10^−4^) with post hoc Tukey HSD test).

Instead, suppressing PM had the surprising effect of increasing the FA rate (p=0.01, paired t-test; **Figure 4G and 4L bottom**). This effect was unique to PM as suppressing V1 decreased FA rate (p=0.02) and suppressing AL or LM had no reliable effect (LM: p=0.94; AL: p=0.71). These changes in hit and FA rate led to a consistent decrease in sensitivity across all areas (d’ for 10% contrast-V1: p=0.009; LM: p<10^−4^; AL: p=0.02; PM: p<10^−3^; paired t-test; **Figure 4H**), suggesting a sensory role for each area. However, while moving the stimulus to the ipsilateral field abolished the increase in threshold in V1, LM and AL (V1: p=0.99, n=3 mice; LM: p=0.24, n=4; AL: p=0.03, n=4; paired t-test; **Figure 4I and Supplementary item 4**), suppression of PM still trended towards increasing the FA rate (p=0.09, **Figure 4J**). Notably, there was no significant difference between contralateral and ipsilateral conditions for mice that had both conditions (p=0.39, n=5 mice, paired t-test), suggesting that the effect of PM on FA rate is not retinotopically-specific. The effect of suppression of PM also did not depend on the genotype used for optogenetic suppression (VGAT/GAD∷Cre (n=4) vs PV∷Cre (n=8): p=0.27; **Figure S3B**), suggesting that the increase in FA rate is not an artifact of the method of suppression.

Thus, we find that sensory integration in V1, LM and AL is required for both orientation discrimination and contrast detection. In comparison, while PM is not required for discriminating orientations, it is involved in performance of the contrast detection task. Namely, while we found suppression of PM weakly increased detection thresholds, the more robust effect was a reliable increase in FA rate, which was independent of visual stimulus input.

### Role of mouse visual areas in go/no-go speed increment detection task

We were surprised that suppressing PM increased FA rate in the contrast detection task but not in the orientation discrimination task. One possible explanation for this discrepancy is that suppressing PM specifically alters computations that are sensitive to noisy changes in neuronal activity. This is a feature of our contrast detection task, where the mouse has to detect small increments in contrast on a noisym background, but not the orientation discrimination task where each high contrast orientation presentation drives a strong signal.

To determine if PM might have a role in regulating noise in the visual system, we next trained the mice to detect a speed increment of a field of randomly moving dots with 0% coherence in stimulus direction (**Figure 5A**). As in the contrast detection task, in the speed increment detection task the mouse has to detect small changes on a noisy background. This task was also structurally similar to the contrast detection task: animals initiated the task by pressing the lever and released the lever within a reaction time window to report a speed increment of the moving dots (0% coherence, baseline speed: 0.5 deg/s, speed increment: 0.94-30 deg/s). We used the same approach to calculate hit rate, detection threshold and FA rate, and compare those between control and area suppression conditions (**Figures 5B-C and Supplementary item 5**). Since neurons in AL and PM have distinct speed preference profiles [18,19,37–40], and therefore may be important for speed change detection, we focused on the role of these two areas in this task.

**Figure 5.**
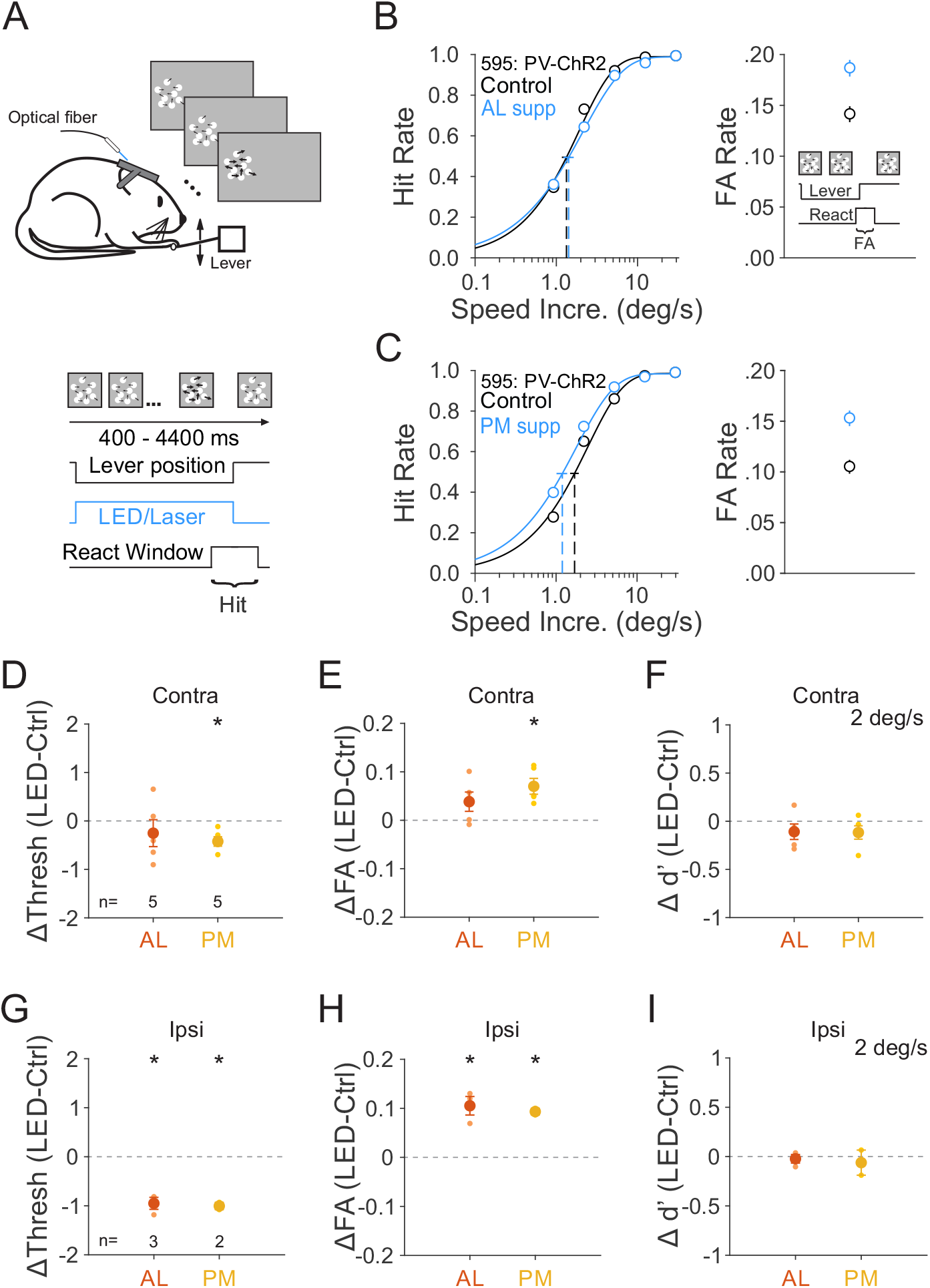
Role of areas AL and PM in detecting speed increment. (**A**) Schematic of speed increment detection task. Base dot speed: 0.5 deg/s, dot coherence: 0%. Thicker arrows indicate an increase in dot speed. (**B-C**) Effect of suppression in area AL (**B**) and PM (**C**) on hit rate (left) and FA rate (right) for an example mouse. Note that this is the same mouse from **Figure 4D-E.** (**D-F**) Summary of effect of suppression in AL and PM in terms of the change in the threshold (**D**), FA rate (**E**) and sensitivity (d’ for 2 deg/s speed increment, **F**). Big circles are the population mean and small circles are data for each mouse. The visual stimulus is presented in the contralateral field of view (relative to the ChR2 injection site). Error bars indicate ± SEM across mice (n=5, 5 for AL and PM). * p<0.05. (**G-I**) Same as **D-F**, for visual stimulus presentation in the ipsilateral field of view. n=3, 2 for AL and PM.

While AL suppression did not have consistent effects on speed increment detection threshold (p=0.42, n=5 mice, paired t-test; **Figures 5D and Supplementary item 5A**) and FA rate (p=0.13, **Figure 5E**), PM suppression consistently decreased detection threshold (p=0.01, n=5 mice) and increased FA rate (p=0.01). In this case, there was no change in sensitivity with PM suppression (p=0.17; **Figure 5F**). Consistent with this lack of a sensory role for PM, these effects were maintained when we presented the visual stimulus ipsilateral to the suppression site (PM-threshold: p=0.05; FA rate: p=0.05; n=2 mice; **Figure 5G-I and Supplementary item 5B**). Thus, as in the contrast detection task, the effects of PM suppression on FA rate in the speed increment detection task were also not specific to the visual input. Interestingly, moving the stimulus to the ipsilateral field also revealed a decrease in threshold and increase in FA rate when suppressing AL (AL-threshold: p=0.02; FA rate: p=0.03; n=3 mice), suggesting that there may be some competing effects of silencing AL during speed increment detection.

In summary, we found that suppression of PM consistently increased FA rate on both the contrast and speed increment detection task independent of effects on sensory, processing, suggesting that PM may have a role in regulation of noise levels in the visual system.

## Discussion

We found functional specialization of mouse HVAs in perception of broadly represented visual features, such as orientation and contrast. While lateral HVAs (LM and AL) contribute to perception of both orientation and contrast, a medial HVA (PM) was not required for discriminating orientations and had a much weaker contribution to contrast detection than lateral HVAs. Notably, suppression of PM consistently increased FA rate in both contrast and speed increment detection tasks independent of visual processing, suggesting a role in regulating noise.

One major finding is that mouse lateral HVAs (LM and AL) are required for perception of simple visual features such as contrast and orientation. Suppression of these areas significantly increases thresholds and decreases sensitivity, and these effects are dependent on the retinotopic alignment of the suppressed neuronal population with the visual stimulus, consistent with a sensory role for these areas in decision-making [33]. This argues that even these relatively simple perceptual tasks require the routing of visual signals through distributed HVAs, and do not rely solely on a pathway from V1 directly to decision-making or motor areas (though see [23]). While the effects of LM and AL on behavior could be the result of “off-target” effects of the removal of excitation to their targets [41,42], we do not think this is the case. For instance, while LM and AL both send excitatory projections back to V1 [43], suppression of these areas do not silence V1 [44], and in some cases actually increase excitability due to a decrease in surround suppression [45]. Further, that suppression of LM and AL have a similar magnitude effect as suppression of V1 suggests that these areas lie downstream of V1 in the perceptual pathway.

The comparable effects of LM and AL on these behaviors also suggests that both of these areas are important for the computation. Given that suppression of either LM or AL only weakly influenced activity in the neighboring area, one would expect asymmetric effects on behavior if only one area were important. One possibility is that these areas have redundant roles, with the intact area partially compensating for the silenced one. Alternatively, the flow of information may be in series from V1➔LM➔AL. Such a pathway is consistent with our anatomical and physiological understanding of the hierarchy [25,46,47], as well as the comparable effects of silencing each of these areas.

Another important finding is the existence of functional specialization for perception of features that are broadly represented across visual areas. In comparison to V1, LM and AL, suppression of PM had no effect on performance of the orientation discrimination task. This is not a trivial result as neurons in PM are similarly responsive to task stimuli, orientation tuned, and suppressed by activation of inhibitory interneurons as the other areas. Moreover, two studies using decoding analysis to classify stimulus orientation from population activity found the decoder performance was either similar between PM, LM and AL [48], or only slightly higher in AL than in LM and PM [17,49]. Thus, our data suggest that there is information present in PM that is not used to perform this orientation discrimination task. This result adds to an accumulating literature suggesting that encoding of task-related information does not always guarantee a causal relationship of that area in behavior [28,50–53].

The third key finding is that roles of HVAs in perceptual decision-making may not be limited to be purely sensory. While there was no effect of PM on the orientation task, suppression of PM did cause a small decrease in sensitivity on the contrast detection task, in part due to an increase in FA rate. This increase in FA rate was independent of the alignment of the suppressed neurons with the visual stimulus, suggesting that the effect is not sensory. However, we do not think that this can be explained by a direct effect on motor output, because we do not see a similar increase in FA rate on the orientation discrimination task. Instead, decreasing activity in PM might disinhibit some target area, thereby leading to an increase in noise. Such an increase in noise might make the downstream decoder more likely to inappropriately pass threshold, especially on a task where the circuit is optimized to detect small increases in activity. Indeed, we observe a similar increase in FA rate when suppressing PM during the speed change detection task, which like the contrast detection task, requires the detection of small changes in spiking at unpredictable times.

The effects of PM on FA rate may act through a variety of targets. PM provides excitatory input to V1, the other HVAs, superior colliculus, higher order thalamic nuclei, and the cingulate cortex, among others [54–56]. These pathways can often have a net inhibitory effect through the recruitment of local inhibition. Indeed, feedback from HVAs to V1 is thought to support some components of surround suppression [45,57,58]. Moreover, a recent study suggests suppression of the lateral posterior (LP) nucleus of thalamus could increase the excitability in V1 resulting in an increase FA rate in a visual discrimination task [59]. Thus, the observed effects of PM suppression in our study could also act through this LP➔V1 circuit. Alternatively, the effects of PM may reflect its feedforward influence on decision-making areas rather than the feedback influence on sensory integration.

Notably, the contribution of PM to regulation of noise in decision-making may not be unique. While suppression of LM and AL do not drive an increase in FA rate during the contrast detection task, nor do they drive a decrease in FA rate as they did in the orientation discrimination task. Thus, this lack of a change may reflect the competition between two mechanisms. Indeed, in the speed increment detection task, we observed an increase in FA and hit rate when suppressing ipsilateral AL without a decrease in FA and hit rate when suppressing contralateral AL. This suggests that the retinotopic-specific visual effects of suppressing an HVA might be obscured by the global non-specific effects. Conversely, the visual effects of suppressing PM during the contrast detection task may be masked to some degree by the changes in false alarm rate.

In summary, although orientation and contrast are broadly represented throughout the mouse visual system, the circuits required for perception of these features are not similarly distributed. While some areas, like LM and AL are clearly required for discriminating orientations and detecting contrast, the role of PM in these tasks is much more subtle and complex. This difference between the medial and lateral HVAs begs the question of what might support the difference in the readout of information. One possibility is the difference in downstream connectivity [55]; another possibility is that the outputs of these areas are differentially integrated by their targets to drive a decision [34,60]. Understanding how each area performs its discrete role will help us understand the computations that link sensation and action.

## Methods

### Animals

All animal procedures conformed to standards set forth by the NIH, and were approved by the IACUC at Duke University. 53 mice (both sexes; 3-30 months old; singly and group housed (1-4 in a cage) under a regular 12-h light/dark cycle; C57/B6J (Jackson Labs #000664) was the primary background with up to 50% CBA/CaJ (Jackson Labs #000654)) were used in this study. *Pvalb-cre* (*tm1(cre)Arbr*, Jackson Labs #008069; n=39; PV∷Cre), *VGAT-ChR2-EYFP* (*Slc32a1-COP4*H134R/EYFP*, Jackson Labs #014548; n=7), *Gad2-IRES-cre* (*Gad2tm2(cre)Zjh*, Jackson Labs #010802; n = 4; GAD∷Cre), Emx1-IRES-Cre (*tm1(cre)Krj*, Jackson Labs # 005628; n=2) and Wild-type (n=1) were used for *in vivo* extracellular electrophysiology (n= 27), and behavior (n=31) experiments. Note five mice were used in both behavior and recording.

### Cranial window implant

Animals were implanted with a titanium headpost and 5 mm cranial window as previously described [61]. Briefly, dexamethasone (3.2 mg/kg, s.c.) and Meloxicam (2.5 mg/kg, s.c.) were administered at least 2 h before surgery. Animals were anesthetized with ketamine (200 mg/kg, i.p.), xylazine (30 mg/kg, i.p.) and isoflurane (1.2-2% in 100% O_2_). Using aseptic technique, a headpost was secured using cyanoacrylate glue and C&B Metabond (Parkell), and a 5 mm craniotomy was made over the left hemisphere (center: 2.8 mm lateral, 0.5 mm anterior to lambda) allowing implantation of a glass window (an 8-mm coverslip bonded to two 5-mm coverslips (Warner no. 1) with refractive index-matched adhesive (Norland no. 71)) using Metabond.

The mice were allowed to recover for one week before habituation to head restraint. Habituation to head restraint increased in duration from 15 min to >2 h over 1-2 weeks. During habituation, imaging and electrophysiology sessions, mice were head restrained while either allowed to freely run on a circular disc (InnoWheel, VWR) or rest in a plastic tube.

### Retinotopic mapping

Retinotopic maps were generated from intrinsic autofluorescence or cortical reflectance (for *VGAT-ChR2-EYFP* mice). For intrinsic autofluorescence (**Figure 1A** and **S1**), the brain was illuminated with blue light (473 nm LED (Thorlabs) or a white light source (EXFO) with a 462 ± 15 nm band pass filter (Edmund Optics)), and emitted light was measured through a green and red filter (500 nm longpass). For cortical reflectance (**Figure S2A**), the brain was illuminated with orange light (530 nm LED (Thorlabs)), and all of the reflected light was collected. For both conditions, images were collected using a CCD camera (Rolera EMC-2, Qimaging) at 2 Hz through a 5x air immersion objective (0.14 numerical aperture (NA), Mitutoyo), using Micromanager acquisition software (NIH). Images were analyzed in ImageJ (NIH) to measure changes in fluorescence (dF/F; with F being the average of all frames) to identify V1 and the HVAs.

To robustly identify the visual areas, we used a two-stage approach for retinotopic mapping (**Figure S1**). First, the monitor was positioned at 45° relative to the body axis, and stimuli were presented at 3 positions (Elevation (El): 10 deg, Azimuth (Az): −10, 10 and 30 deg, 40° sinusoidal gratings, drifting at 2 Hz, 10 s duration, 10 s inter-trial interval) to activate locations in the contralateral visual field (**Figure S1A**). This allowed us to identify sites of retinotopic reversals to define area identity and boundaries [7,16,20]. Next, we positioned the monitor at 0° relative to the body axis, and stimuli were presented at the location used during the behavioral task (El:10, Az: 30-40 deg, green in **Figure S1B**). We used these maps to measure the distance between areas (**Figure 1D-E**) and target our viral injections to the retinotopic location activated by the task stimulus. Vascular landmarks were used to identify targeted sites for targeted viral injections and electrophysiology. In some cases, PM and AM were not entirely separable in the location of the behavior task. But we could always easily define their boundaries by using more lateral stimuli positions (**Figure S1A**).

### Viral injection

We targeted V1, or AL and PM, or LM and PM in PV∷Cre or GAD∷Cre mice for viral expression of Channelrhodopsin2 (ChR2) or Chronos. Dexamethasone (3.2 mg/kg, s.c.) was administered at least 2 h before surgery and animals were anesthetized with isoflurane (1.2-2% in 100% O_2_). The coverslip was sterilized with 70% ethanol and the cranial window removed. A glass micropipette was filled with virus (AAV5.EF1.dFloxed.hChR2.YFP (titer: 3.74e12 GC/ml; UPenn CS0384), AAV9.CAGGS.FLEX.ChR2.tdTomato (titer: 2.44e12 GC/ml; Addgene 18917) or AAV1.Syn.FLEX.Chronos.GFP (titer: 4.80e12 GC/ml; Addgene 62722)), mounted on a Hamilton syringe, and lowered into the brain. 30-50 nL of virus (30 nl for HVAs; 50 nl for V1) were injected at 250 and 500 μm below the pia (30 nL/min); the pipette was left in the brain for an additional 10 minutes to allow the virus to infuse into the tissue. Following injection, a new coverslip was sealed in place, and for behavioral experiments, an optical fiber (400 μm diameter; Doric Lenses) was attached to the cranial window above the injection site. Optogenetic behavioral experiments and electrophysiology experiments were conducted at least two weeks following injection to allow for sufficient expression. On average, the expression covered 0.77±0.28, 0.38±0.09, 0.52±0.12, and 0.33±0.03 mm^2^ for V1 (n=5), LM (n=7), AL (n=12) and PM (n=15) respectively (p=0.07, one-way ANOVA across areas).

### Visual stimulation

Visual stimuli were presented either on a 144-Hz (Asus) or 120-Hz (Samsung) LCD monitor for electrophysiology and behavior experiments, respectively. Monitors were calibrated with an i1 Display Pro (X-rite) for mean luminance at 50 cd/m^2^ and positioned 21 cm from the eye. The stimulus presentation protocols in behavior and electrophysiology experiments are described in each section.

### Behavioral task

Animals were water scheduled and trained to discriminate the orientation of visual stimuli, or detect the appearance of a visual stimulus with varying contrast, or detect the increment of speed of the moving dots by manipulating a lever. The behavior training and testing occurred during the light cycle. All behavioral control and stimulus presentation used MWorks (http://mworks-project.org), and custom software in MATLAB (MathWorks).

The go/no-go orientation discrimination task was trained and performed as previously described [34]. Briefly, each trial was initiated when the ITI (3s) had elapsed and the mouse had pressed the lever. Trial start triggered the presentation of a series of 100 ms static sinusoidal, gabor patches (diameter: 30°, spatial frequency (SF): 0.1 cycle/deg, contrast: 100%, positioned at an eccentricity of 30° - 40° in azimuth and 0° - 10° in elevation) followed by a target orientation of the same parameters but of a different orientation. The spatial frequency value was chosen to ensure the task stimuli drive all HVAs equally well (**Figure 3C**). The target orientation occurred with a variable delay (flat distribution) after at least two distractor presentations (up to 10 distractors). Following each target, additional distractors were presented until the mouse either released the lever or the reaction time expired. Within each trial, each stimulus presentation (distractors and target) was separated by a mean-luminance ISI (either constant at 250 ms, or randomized between 250, 500 and 750 ms). Each trial had the possibility of having a target presentation, if the mouse held the lever through the all of the preceding distractor presentations. Mice received water reward only if they released the lever within 100-650 ms (sometimes extended to 1000 ms) after a target occurred. If mice released the lever before reaction time began (early release) or failed to release the lever by the time the reaction time expired (miss), the trial would be aborted and additional time (2-4s) would be added to the ITI.

The go/no-go contrast detection task was trained and performed as previously described [36]. Briefly, as in the orientation discrimination task, each trial was initiated when the ITI (3s) had elapsed and the mouse had pressed the lever. Following a variable period (from 700-3500 ms), a static sinusoidal target grating (diameter: 30°; SF: 0.1 cycle/deg; azimuth of 30° - 40°; elevation of 0° - 10°) of a variable contrast (range: 4%-100% contrast) was presented. Delays after errors were also added to discourage lapses and early releases.

For the go/no-go speed increment detection task, we first trained mice to detect an increase in speed of the random dot kinetogram (speed increase from stationary to 30 deg/s; full-field; 100% coherence in nasal to temporal direction; dot density: 0.025; dot size: 3 deg) that started at the end of the required hold time (400 ms). Once the animals started to respond to the change of speed by releasing the lever, we gradually made the task harder by: 1) decreasing the dots coherence to 0%; 2) increasing the random delay (final max value: 4 s) between the lever press and speed increment; 3) shrinking the field size and moving it to more eccentric positions (30° in diameter; azimuth of 35°; elevation of 10°); 4) increasing baseline speed from 0 to 0.5 deg/s; and 5) introducing hard target speeds (difference from the baseline speed: 0.94-30 deg/s) to probe speed increment detection threshold. Delays after errors were also added to discourage lapses and early releases.

We delivered blue light to the brain though an optic fiber from a 473 nm LED (Thorlabs) or a 450 nm laser (Optoengine) and calibrated the total light intensity out of the fiber. For all behavior tasks, the light was delivered on 50% of trials (randomized with control trials) for the entire duration of the trial and terminated after either animal released the lever or the reaction window expired. A black cap was used to block the blue light to minimize the chance of animal detecting the light delivery. Note that for behavior tests using VGAT-ChR2 transgenic mouse line, the light power never exceeded 0.4 mW to minimize spatial spread of cortical suppression (**Figure S3**, [31]).

### Extracellular electrophysiology

Electrophysiological signals were acquired with a 32-site polytrode acute probe (A4×8-5mm-100-400-177-A32 (4 shanks at 400 μm spacing, 8 site/shank at 100 μm spacing, NeuroNexus) through an A32-OM32 adaptor connected to a Cereplex digital headstage (Blackrock Microsystems). Unfiltered signals were digitized at 30 kHz at the headstage and recorded by a Cerebus multichannel data acquisition system (Blackrock Microsystems).

On the day of recording, the cranial window (and the optic fiber, if it was already implanted from behavioral experiments) was removed, and a small durotomy performed to allow insertion of the electrode in visual cortex. A ground wire was connected via a gold pin cemented in a burrhole in the anterior portion of the brain. The probe was slowly lowered into the brain (over the course of 15 min with travel length of around 800 μm) until the most superficial recording site was in the brain and allowed to stabilize for 45 - 60 min before beginning recordings. The 4-shank probe was targeted so that one shank was centered on the injection site. In some cases, all four shanks were in V1; in other cases, they might span across V1 and HVAs, or all be in lateral or medial HVAs.

For optogenetic stimulation, the optic fiber was held in place via an articulated arm (Flexbar, SKU: 14830) to allow light delivery (473 nm LED, Thorlabs) to the injection site. While mice were viewing gray screen (no visual stimuli), constant blue light was delivered with a duration of either 3.5 s or 3.8 s, which was randomly interleaved with control trials with no light delivery. Between each trial, there was a 4 s ITI to match the condition as in the behavioral task. For viral injections, the light was delivered through 400 μm optic fiber (Thorlabs) with the mean light power 0.34±0.1 mW (range: 0.1-1.5 mW). For transgenic VGAT-ChR2 mice, the light was delivered through 50 μm optic fiber (Thorlabs) and a collimator (Edmund Optics) with the mean light power 0.42±0.01 mW (range: 0.4-0.5 mW). The light delivery methods and light power matched those in the behavioral experiments (**Figures 2L**, **3L** and **S3**).

To measure orientation tuning and responses to task stimuli in V1 and the HVAs (**Figure 3B-C**), we presented five repetitive static, 100 ms sine-wave gratings with the same orientation (randomized from 0°, 30°, 60°, 90°, 120°, 150°) with an ISI of either 250 or 500 ms and an ITI of 4 s. The contrast of all the stimuli was 100% and spatial frequency was 0.1 cycle/degree, matching those used in the orientation discrimination task. Thus, we used the response to the first stimulus in each trial to measure each neuron’s orientation tuning; only trials with an orientation of 0° were used to measure responses to the task distractor stimulus.

### Data processing

#### Behavior processing and analysis

All behavioral processing and analysis were performed in MATLAB. All trials were categorized as either an early release, hit, or miss based on the time of release relative to target onset: responses occurring earlier than 100 ms after the target stimulus were considered early releases; responses occurring within the reaction window (orientation/contrast task: 200-550 ms; speed task:200-700 ms) after the target were considered hits; failures to respond before 550 ms (or 700 ms for speed task) after the target were considered misses. For the orientation task, the same reaction window was used following each distractor to calculate false alarm (FA) rate (FAs are a subset of early releases). Thus, each distractor presentation was categorized as either a FA or correct reject (CR), and each target presentation was either a hit or a miss. Since there were no distractors presented in the contrast and speed increment detection task, we calculated FA rate by simulating the timing of potential distractor presentations to match the distribution of target presentations and assessing the probability of the mouse releasing the lever during these windows.

Behavioral sessions were manually cropped to include only consecutive trials in each session with stable periods of performance. Sessions were selected based on the following criteria: 1) at least 40% of trials were hits; and 2) less than 50% of trials were early releases. Based on these criteria, the data in orientation discrimination (**Figure 2**) included 17 ± 2 (range: 6-48) sessions for each mouse with 5675 ± 596 trials (range: 1982-14851) for contralateral condition; 17 ± 2 (range: 7-36) sessions for each mouse with 5046 ± 698 trials (range: 975-10532) for ipsilateral controls; the data in contrast detection (**Figure 4**) included 19 ± 2 (range: 5-62) sessions for each mouse with 5938 ± 879 trials (range: 1796-25302) for contralateral condition; 15 ± 2 (range: 4-30) sessions for each mouse with 4291 ± 634 trials (range: 1281-10344) for ipsilateral controls; the data in speed increment detection (**Figure 5**) included 32±2 sessions (range: 22-44) for each mouse with an average of 7091 ± 795 trials per mouse (range: 5028-13658) for contralateral condition; 20 ± 4 (range: 10-32) sessions for each mouse with 3945 ± 806 trials (range: 1433-6455) for ipsilateral controls.

Hit rate was computed from the number of hits and misses for each stimulus type:

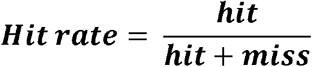

FA rate was computed from the total number of FAs and CRs in the session:

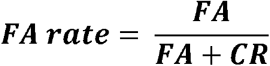

Signal detection theory [62] was applied to measure neuronal sensitivity (d’). Extreme values of hit and FA rate (i.e. 0 and 1) were replaced with 0.5/n and (n-0.5)/n, respectively, where n is the number of target or distractor trials[35,63]. d’ was then calculated as follows:

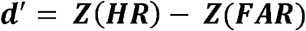

where Z is the inverse of the cumulative distribution function of the normal Gaussian distribution; HR is hit rate and FAR is FA rate.

Since threshold varied across mice and not all the mice were tested at exactly the same orientation, contrast, or dot speed, we chose the stimuli values that were near the threshold to summarize d’ across animals. Thus, the hit rate for 22.5° orientation, 10% contrast, 2 deg/s dots speed were extrapolated based on a Weibull function fitted from the psychometric curve for each mouse.

For characterizing the dependence of performance across trial length for the orientation discrimination task (**Figure S4B-C**), we divided trials into four bins (950-1450, 1450-1950, 1950-2450, 2450-8000 ms) and only included mice that had at least 20 trials for each target orientation at each bin to ensure reliable measures. Thus, 5/6, 4/7, 5/7, 5/5 mice were selected for V1, LM, AL and PM respectively.

#### Electrophysiology processing and analysis

Individual single units were isolated using the SpyKing CIRCUS package (http://spyking-circus.readthedocs.io/en/latest/). Raw data were first high pass filtered (> 500 Hz) and spikes were detected when a filtered voltage trace crossed threshold (9-13 median absolute deviations computed on each channel). A combination of density-based clustering and template matching algorithms were used to automatically cluster the spikes. The resulting clusters were then inspected and adjusted manually using a MATLAB GUI. Clusters with refractory period violations (< 2 ms, >1% violation) in the auto-correlogram and that were not stable across the whole recording session were discarded from the dataset. Clusters were combined if they met each of three criteria by inspection: 1) similar waveforms; 2) coordinated refractory periods in the cross-correlogram; 3) similar inter-spike interval distribution shape. Since each contact site is at least 100 μm apart, unit position was assigned to the contact site which exerts the biggest waveform.

To quantify effects of inhibition by exciting interneurons, spike times across trials (>30 trials) were first converted to peri-stimulus time histograms (PSTHs, bin size: 100 ms; align to the onset the light delivery, with 2 s baseline period). We used a paired t-test to exclude neurons that were significantly driven by blue light, presumably interneurons, from later analysis. To determine the inhibition efficacy, we measured the normalized suppression as:

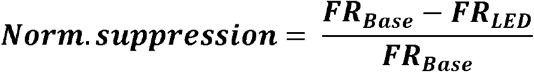

where FR_LED_ is the mean firing rate during the light delivery, FR_Base_ is the mean baseline firing rate. Thus, value of 1 means entirely suppressed; values smaller than 0 suggests excited by light. We then averaged these values across neurons to obtain spatial resolution profiles across cortical depth in the light center (**Figure S2C**) and cortical distance away from light center (**Figures 1C and S1D**). Note that for both of the suppression methods, the degree of suppression was stable across cortical depth in the light center (viral: p=0.11, n=73 cells; transgenic: p=0.63, n=53 cells; Kruskal-Wallis test across depths, excluding ChR+ cells).

To quantify the percentage of area coverage of suppression, we measured the spatial decay constant as a single exponential fit to the response as a function of tangential distance from the injection center, assuming a homogeneous decay constant across visual cortex for a given optogenetic method (**Figure 1**, PV/GAD:ChR2 (viral injection); **Figure S2**, VGAT-ChR2 transgenic mouse line). We then estimated the area of activation within each visual area, and the relative distance between each, using the intrinsic imaging (**Figure 1A** left and **S1B**) obtained in response to stimuli with position matched to the behavior tasks (elevation 10°, azimuth 30-40°, but with stimulus size larger than used in the behavior: 40° here versus 30° in the behavior). For each mouse, the active region in each area was fitted with an oval to identify its center and area (**Figure 1D**). Finally, we approximated the percentage of suppression coverage within and across areas by calculating the fraction of each visual cortical area encompassed by the spatial decay constant centered on each area. We then averaged across mice (n=15) to obtain the inhibition coverage profiles traversing through six visual areas (**Figure 1E and S2E**, values were binned as 0-0.1%, 0.1-2%, 2-50%, 50-98%, 98-100%).

Visually evoked responses of each unit were measured based on average peri-stimulus time histograms (PSTHs, bin size: 20 ms) over repeated presentations (>25 trials) of the same stimulus. Response amplitudes were measured by subtracting the firing rate at the time of the visual stimulus onset from the value at the peak of the average PSTH within a window of 0-130 ms after the visual onset. For further analysis, only “responsive cells” that had statistically significant averaged visually evoked responses during 0-130 ms window after first visual presentation onset using a paired t-test comparing to the baseline response (average response during 0-130 ms window before visual onset). To determine how well the stimulus orientation information is encoded in each area, we measured circular variance [64] for each “responsive cell” (significantly responsive to collapsed all orientations):

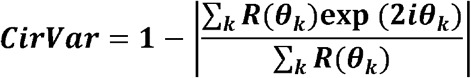

where R(θ_k_) is the response to a sampled orientation θ_k_ (choosing from 0°, 30°, 60°, 90°, 120°, 150°) and i is the imagery unit. Thus, CirVar has the range from 0 to 1, with value of 0 indicating maximal orientation selectivity and value of 1 indicating lack of orientation selectivity.

### Experimental Design and Statistical Analysis

All behavioral and neuronal data were tested for normality using a Lilliefors test. While behavioral measures were normally distributed, electrophysiological measures of spike rates were not. Therefore, behavioral data were compared with either a t-test or ANOVA with post hoc Tukey HSD test for datasets with two and multiple groups, respectively. However, for the neuronal activity we used only non-parametric tests (Wilcoxon signed rank test and Friedman test with post hoc Tukey HSD test to compare two and multiple groups, respectively). Sample sizes were not predetermined by statistical methods, but are similar to other studies. The numbers of cells, animals or experiments were provided in the corresponding text, figures and figure legends. All error values in the text are SEMs unless otherwise specified. Data collection and analysis were not performed blind to experimental conditions, but all visual presentation and optogenetic stimulation conditions in behavior and electrophysiology experiments are randomized.

## Data and code availability

All relevant data and code are available from the corresponding author upon reasonable request.

## Acknowledgements

We thank J. Sims, A. McKinney and B. Gincley for assistance with behavioral training; E. Burke, K. Leonard, M. Fowler, J. Isaac and K. Murgas for surgical assistance; Z. Xu for assistance with software development; G. Field, C. Hull, S. Lisberger, F. Wang and members of the Hull and Glickfeld labs for helpful discussions and comments on the manuscript. This work was supported by an NIH Director’s New Innovator Award (DP2-EY025439), the Pew Biomedical Trusts, and the Alfred P. Sloan Foundation (L.L.G).

## Author Contributions

M.J. and L.L.G. designed the experiments. M.J. collected and analyzed the electrophysiology and behavior data. M.J. and L.L.G wrote the manuscript.

---

**Figure S1.**
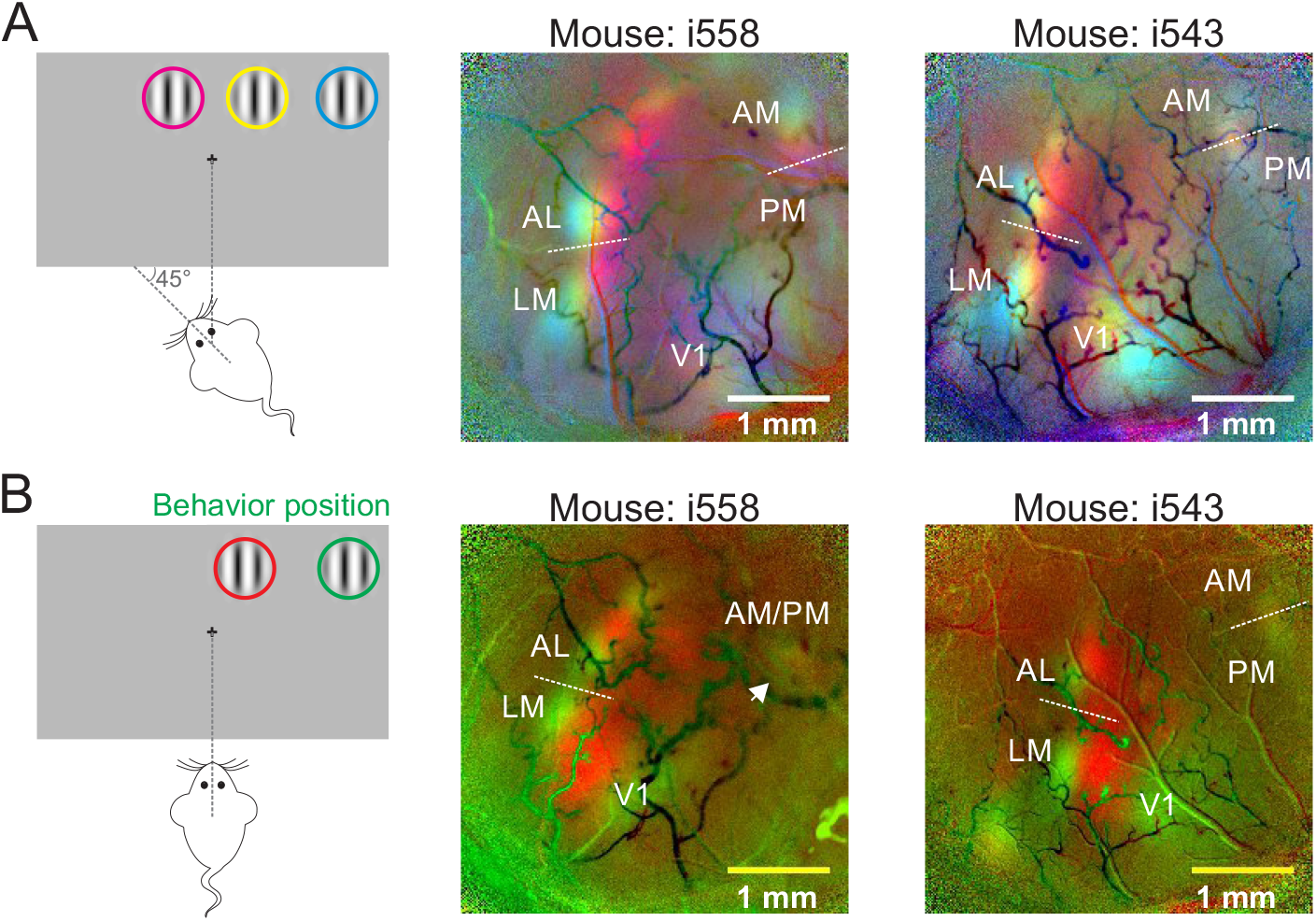
Identify boundaries between visual cortical areas using intrinsic autofluorescence imaging. (**A**) Change in intrinsic autofluorescence for two example mice (right) in response to visual stimuli presented at three positions (left) when animal is rotated 45° relative to the screen. Colors in the map match the positions outlined in the schematic (azimuth −10°, 10° and 30°, elevation of 10°). Lines show identified boundaries between AL and LM, AM and PM. (**B**) Same as **A**, for responses to the two positions (azimuth 10-20° and 30-40°, elevation 10°) when the same animals are perpendicular to the monitor. Arrowhead points to the place where AM and PM are not entirely separable at the behavior position (green). Note that mouse i558 is the same mouse in **Figure 1A**.

**Figure S2.**
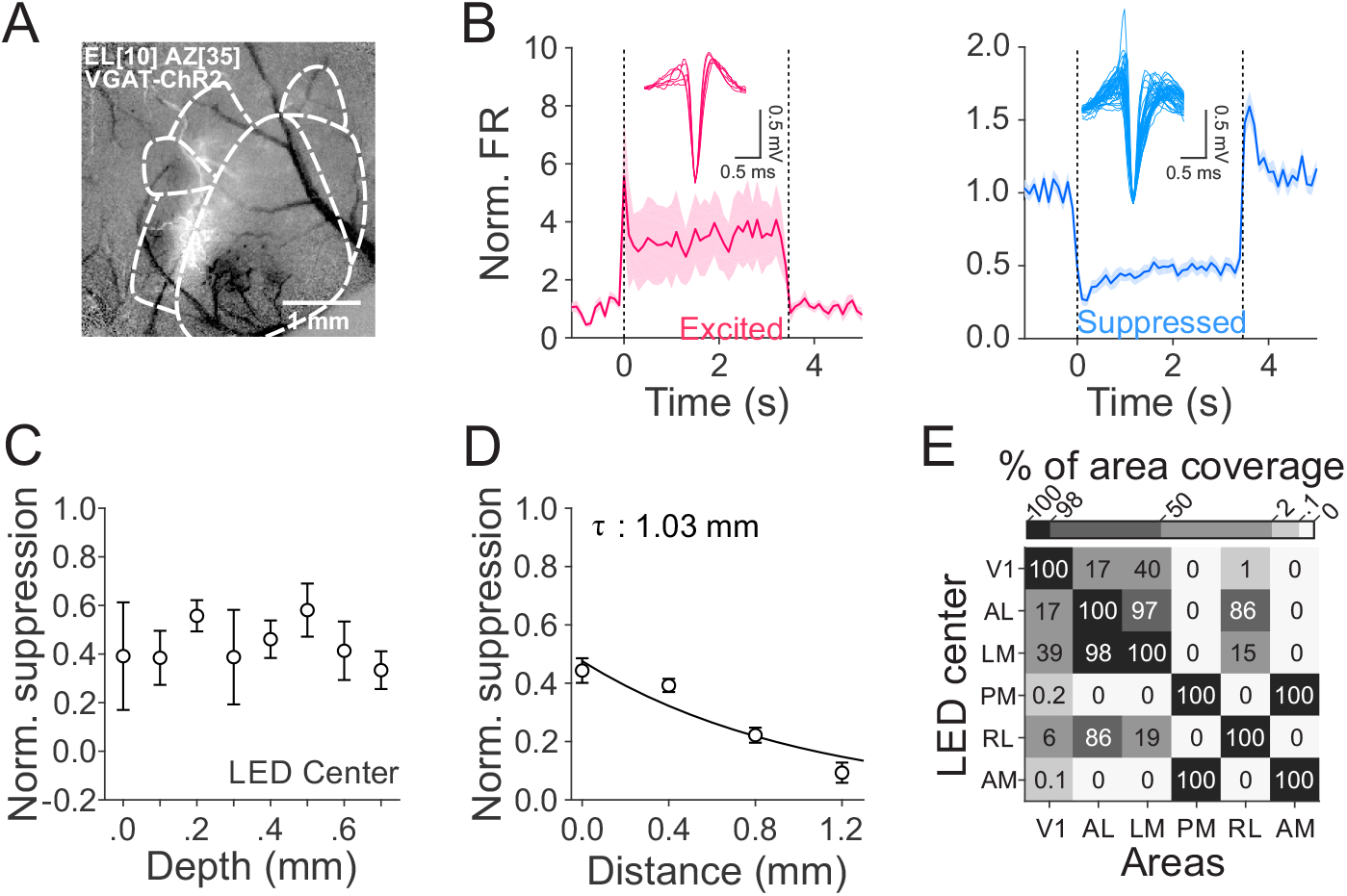
Efficacy and spatial resolution of optogenetic inhibition in VGAT-ChR2 transgenic mouse line. (**A**) Cortical reflectance in response to a visual stimulus at elevation 10°, azimuth 35°, size 40°. Decreases in reflectance reflect activation. (**B**) Normalized spontaneous firing rates (FR) of cells that are excited (left, red, n=6 cells) and suppressed (right, blue, n=42 cells) when stimulated with 450 nm laser. Dashed lines indicate laser onset and offset. Insets are the waveforms of single units. Light power: 0.4-0.5 mW across 9 experiments (n=2 mice). (**C**) Normalized suppression as a function of cortical depth in the laser center. Error bars are SEM across cells (n=5, 3, 5, 6, 11, 8, 7, 8 cells, from superficial to deep). (**D**) Normalized suppression as a function of distance from the laser center (0 mm). Error bars are SEM across cells (n=53, 125, 64, 20, from 0 to 1.2 mm). The decay constant (τ) was calculated via a single exponential fit. (**E**) Mean percentage of area coverage of inhibition across mice (n=15).

**Figure S3.**
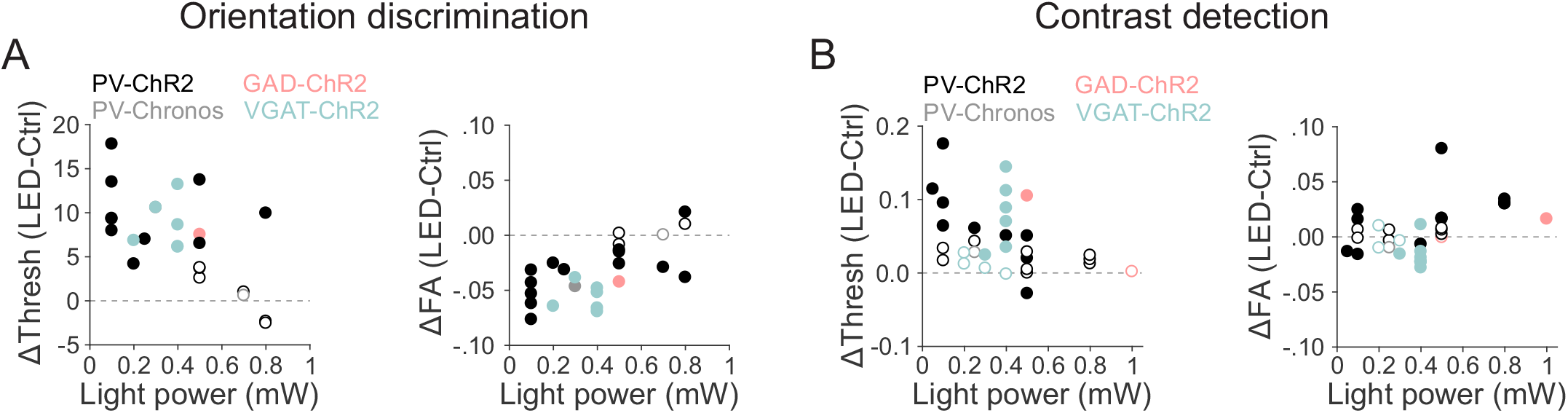
Effects of suppression method and light power on behavior performance in orientation discrimination and contrast detection tasks. (**A**) Effects of area suppression on orientation discrimination threshold (left) and FA rate (right) as a function of light power. Different colors denote different suppression methods. Filled and open circles denote significant and non-significant difference between control and LED trials. (**B**) Same as **A**, for the performance in the contrast detection task.

**Figure S4.**
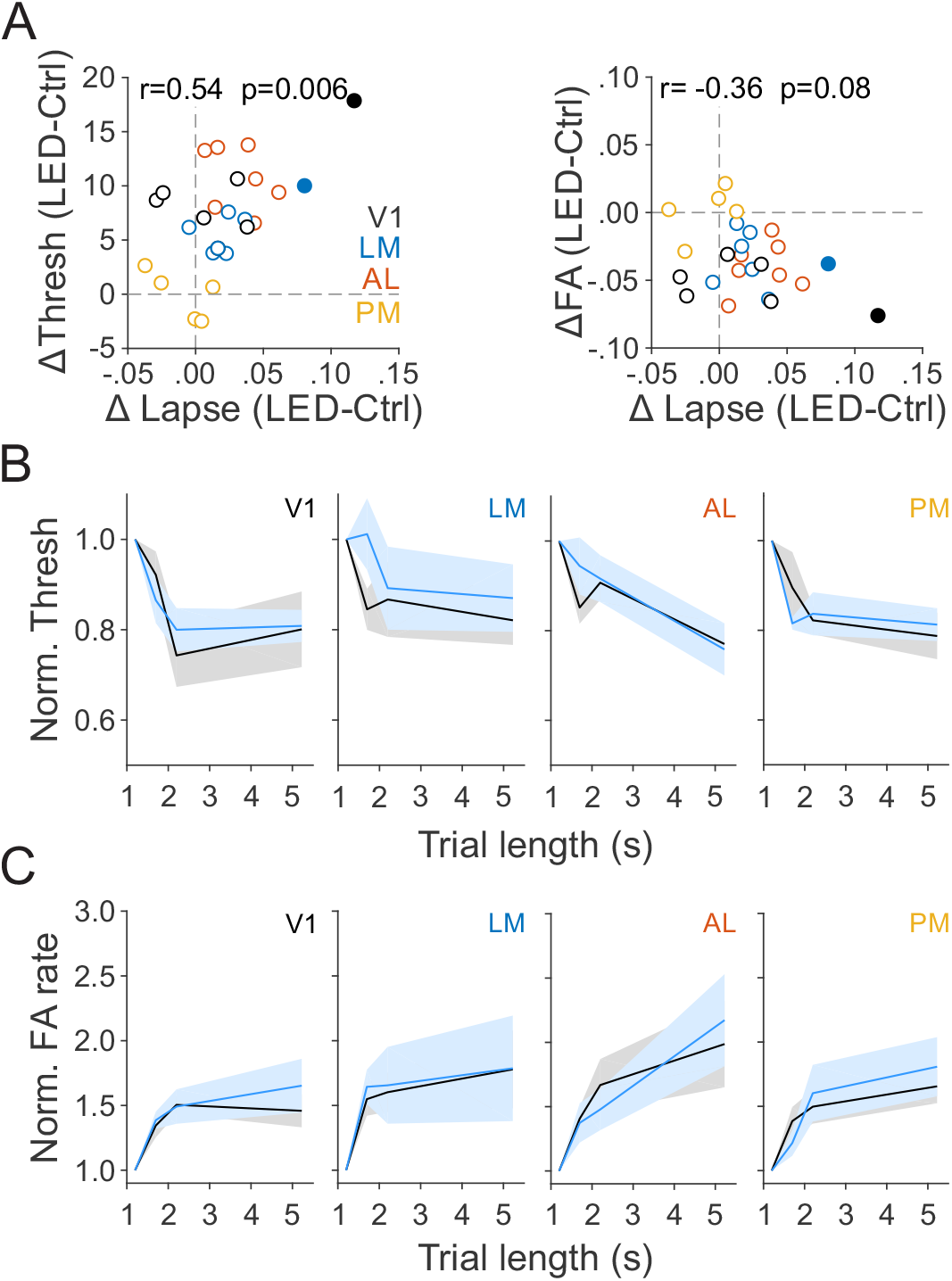
Lack of effects of area suppression on engagement and expectation, related to Figure 2. (**A**) Changes in orientation discrimination threshold (left) and FA rate (right) plotted against changes in lapse rate. Different colors denote different visual areas. Filled and open circles denote significant and non-significant difference in lapse rate between control and LED trials. (**B**) Summary of normalized orientation discrimination threshold over trial length bins (950-1450, 1450-1950, 1950-2450, 2450-8000 ms) for control (black) and LED (blue) trials. Shaded areas are SEM across mice (n=5, 4, 5, 5 for V1, LM, AL, and PM respectively). (**C**) Same as (**B**) for FA rate.

**Supplementary item 1.**
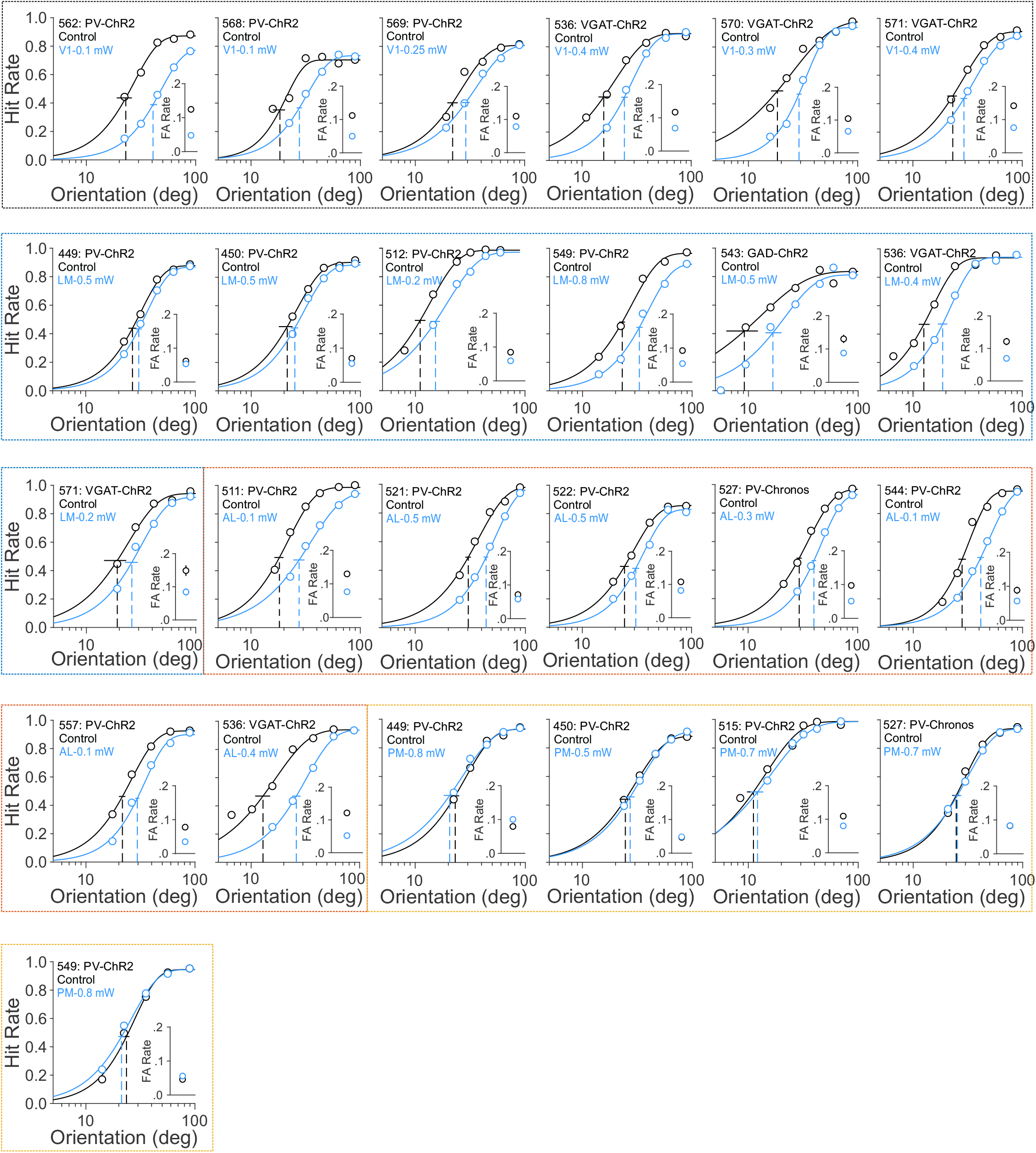
Behavior performance of all mice for discriminating orientations when visual stimuli were in the contralateral field of view (relative to the suppression site), related to Figure 2. Figures are organized by areas (colored outlines: black- V1; blue- LM; red-A L; yellow- PM).

**Supplementary item 2.**
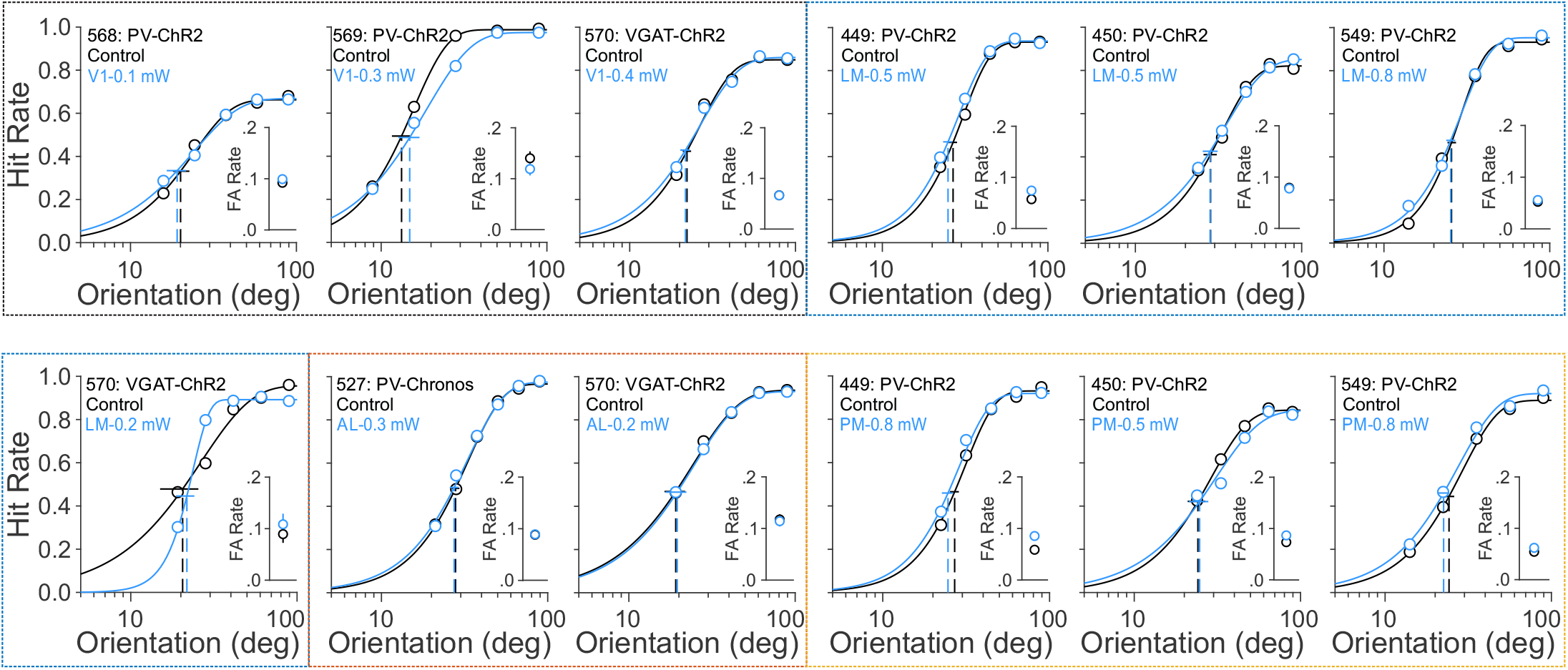
Behavior performance of all mice for discriminating orientations when visual stimuli were in the ipsilateral field of view (relative to the suppression site), related to Figure 2. Figures are organized by areas (colored outlines: black- V1; blue- LM; red- AL; yellow- PM).

**Supplementary item 3.**
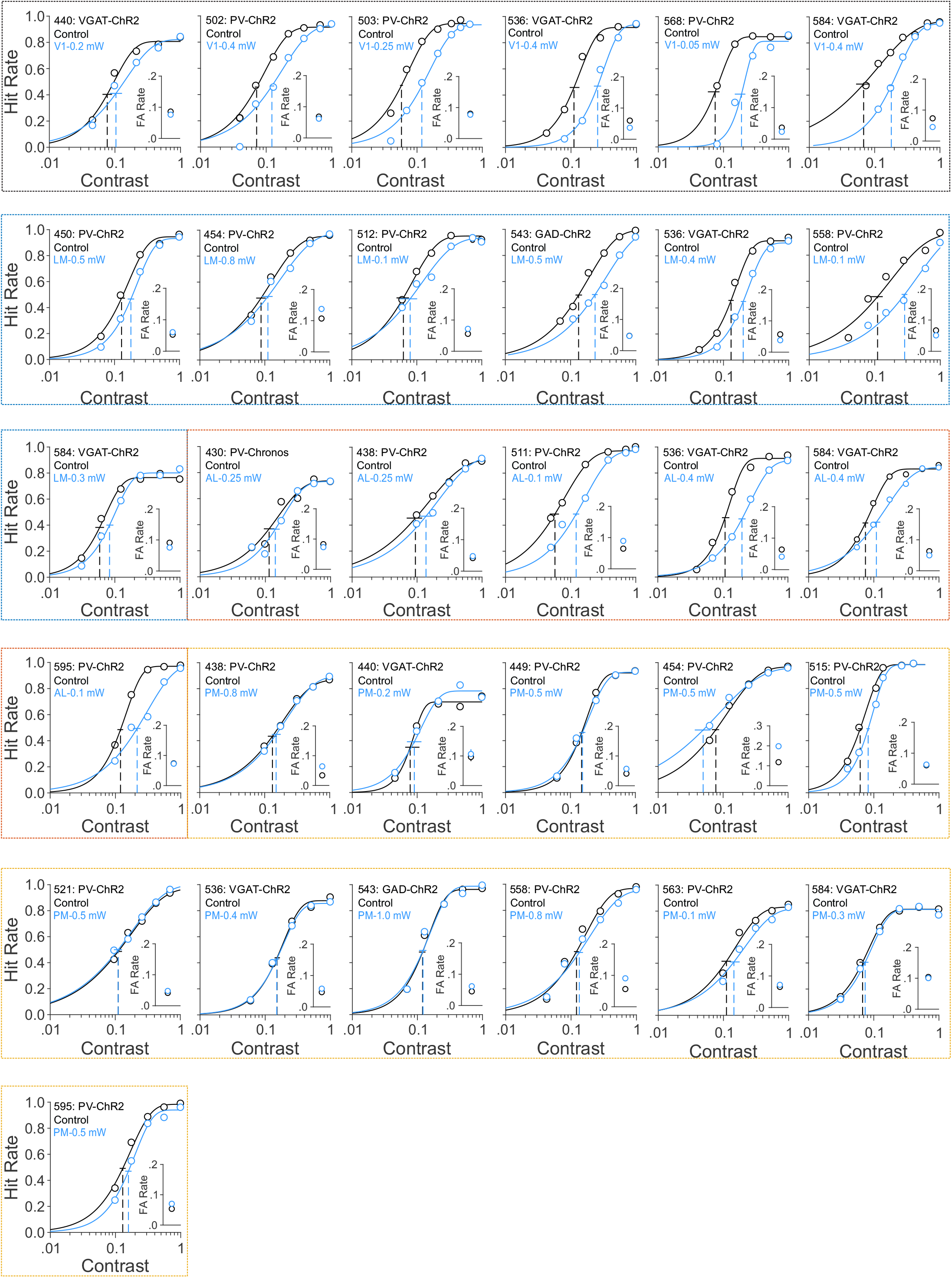
Behavior performance of all mice for detecting contrast when visual stimuli were in the contralateral field of view (relative to the suppression site), related to Figure 4. Figures are organized by areas (colored outlines: black- V1; blue-LM; red- AL; yellow- PM).

**Supplementary item 4.**
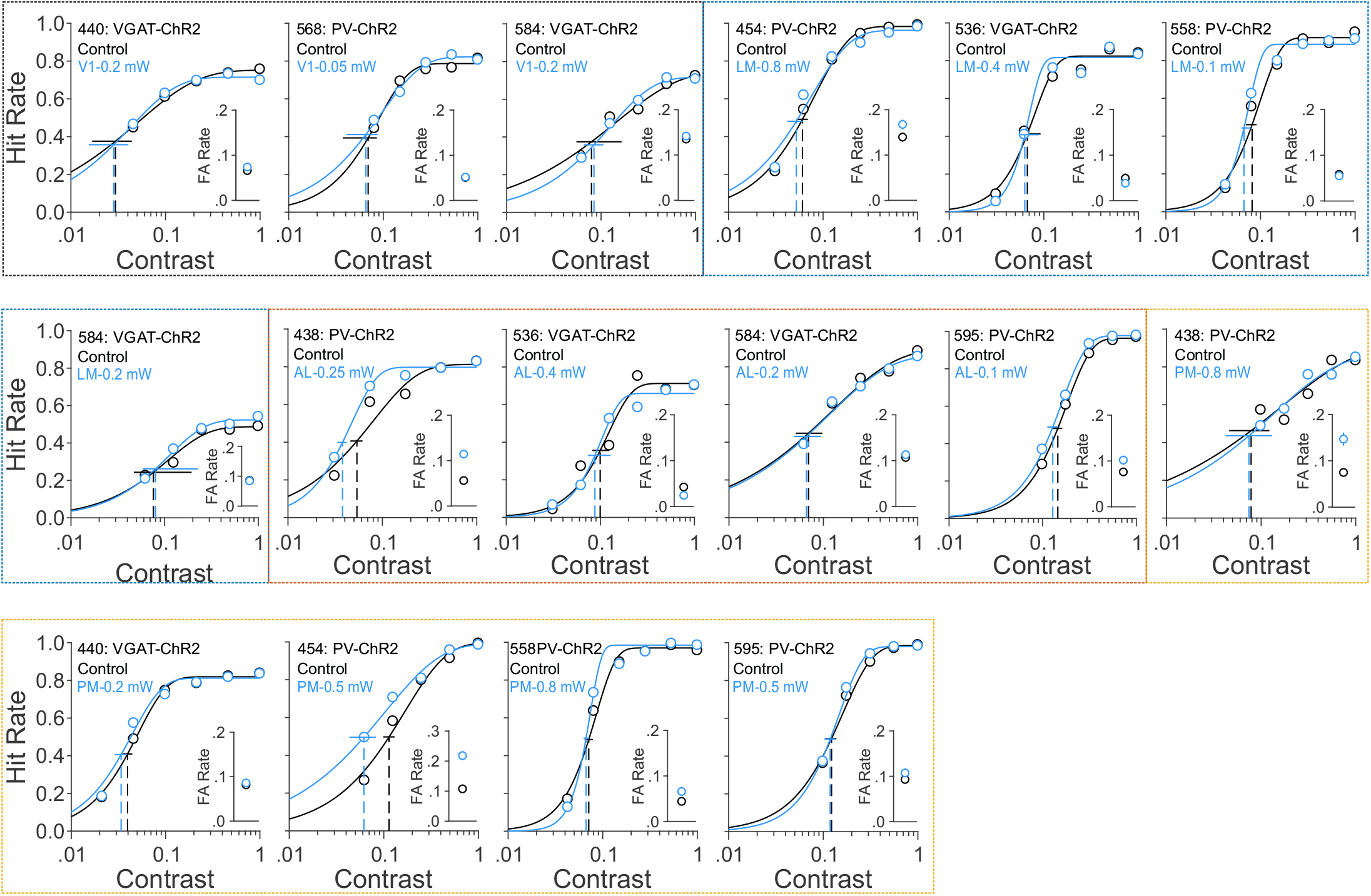
Behavior performance of all mice for detecting contrast when visual stimuli were in the ipsilateral field of view (relative to the suppression site), related to Figure 4. Figures are organized by areas (colored outlines: black- V1; blue- LM; red- AL; yellow- PM).

**Supplementary item 5.**
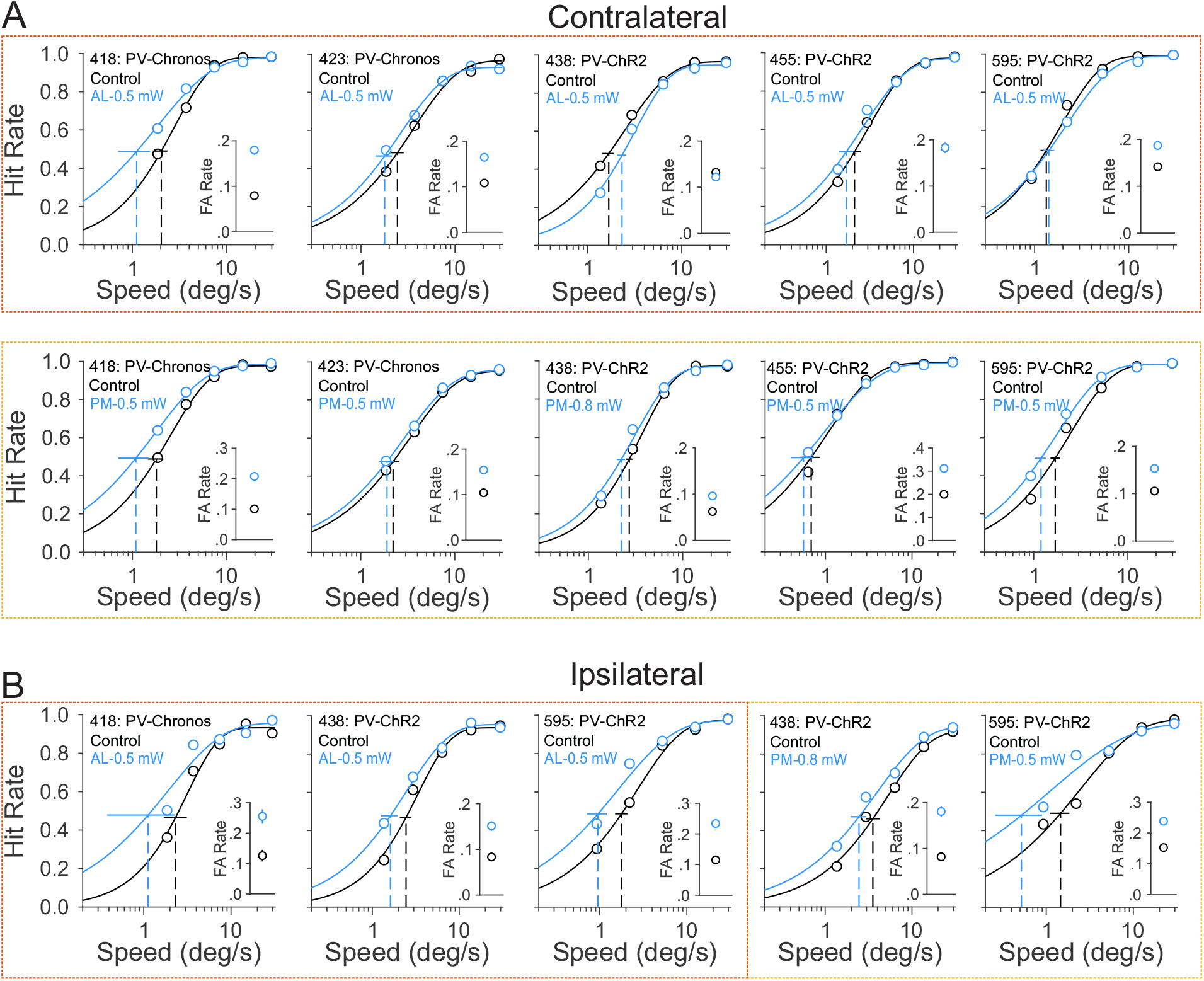
Behavior performance of all mice for detecting speed increment when visual stimuli were in the contralateral (A) and ipsilateral (B) field of view (relative to the suppression site), related to Figure 5. Figures are organized by areas (colored outlines: black- V1; blue- LM; red- AL; yellow- PM).

